# The Primate Hippocampus Constructs a Temporal Scaffold Anchored to Behavioral Events

**DOI:** 10.1101/2025.11.12.687961

**Authors:** Jon W Rueckemann, Yoni Browning, Autumn J Mallory, Brian Kim, Adrienne L Fairhall, Elizabeth A Buffalo

## Abstract

The hippocampus has been attributed a range of divergent functions, including roles in memory^1,2^ and navigation^3–5^, but how its moment-to-moment neuronal activity supports cognition remains poorly understood. The activity of individual hippocampal neurons is correlated with numerous perceptual features and task variables^6–14^, raising the question of whether these response properties reflect distinct mechanisms or support a single generalized computation. Here, we show that these diverse response properties reflect a unified organizing principle in which the hippocampus segments experience into discrete events, with population activity transitioning between discrete neural states at behaviorally salient moments. Recording from monkeys performing a virtual spatial alternation task, we found that population activity did not evolve smoothly over time but instead shifted abruptly at each relevant event. These discontinuities segmented activity into distinct ensembles, effectively chunking separate task epochs. Notably, many neuronal responses persisted across visually distinct environments, demonstrating that these dynamics reflect abstract task structure rather than specific sensory features. These results reveal that the hippocampus constructs a temporal scaffold anchored to relevant behavioral events, with each neural state tracking a distinct task phase. This organizational principle may explain the diverse neural correlates observed across studies: rather than individually encoding perceptual or behavioral variables, hippocampal neurons collectively signal the current phase of a behavioral sequence. Our findings suggest that the hippocampus parses experience into meaningful elements and tracks “position” within a learned behavioral structure.

## Introduction

Damage to the hippocampus impairs memory^15–17^ and cognition^18–21^, yet we lack a fundamental account of what this structure computes. Much of what we know about its physiology comes from rodents navigating space, where neurons fire at specific locations and have been interpreted as encoding a map of the environment^3–5^. However, studies have also shown that hippocampal neurons respond to tones^6^ and odors^9^, track elapsed time during delays^11,22,23^, and modulate their activity by behavioral context^12,14,24,25^, revealing that the hippocampus encodes dimensions other than physical location. This diversity of responses has supported views in which individual hippocampal neurons act as conjunctive feature detectors that bind sensory, temporal, and contextual information within each moment^26,27^.

A complementary perspective shifts the focus from representing individual moments to capturing the continuity that links them across time^28–37^. In this view, the hippocampus functions as a sequence generator whose activity evolves systematically across events^38^, producing structured patterns that bridge the transitions of experience. In support of this view, studies have shown that hippocampal activity bridges delay periods^39^, predicts upcoming actions^24,40–42^, tracks iterations across trials^43^, reflects ordinal relationships^44,45^, and links events into a holistic sequence^46–53^, indicating that the hippocampus conveys not only what is happening but how the current moment is embedded within the broader temporal progression of behavior. At the same time, hippocampal responses mark behaviorally-defining moments, as evidenced by acute changes in hippocampal activity at boundary events in humans^51,54–60^ and by preferential engagement of single units at reward^8,61–66^ and other task anchors^61,64–67^ in rodents. These perspectives converge on the idea that the hippocampus does more than encode discrete features—it tracks the evolving organization of experience, integrating the continuity of ongoing activity with boundaries sculpted both by external events and the organism’s own interpretation of transitions.

## Results

To examine how these principles manifest at the neuronal level in primates, we recorded neuronal activity from the hippocampus of three adult male rhesus monkeys (*Macaca mulatta*) as they performed a spatial alternation task in an immersive virtual environment (Fig. 1a). Subjects used a joystick to navigate Y-shaped mazes in which they were rewarded for alternating between left and right arms across successive trials (Fig. 1b; Supp Video 1). Subjects performed the task in multiple, visually distinct Y-shaped mazes within a recording day, allowing us to dissociate responses tied to specific sensory features from those reflecting task structure.

**Figure 1.**
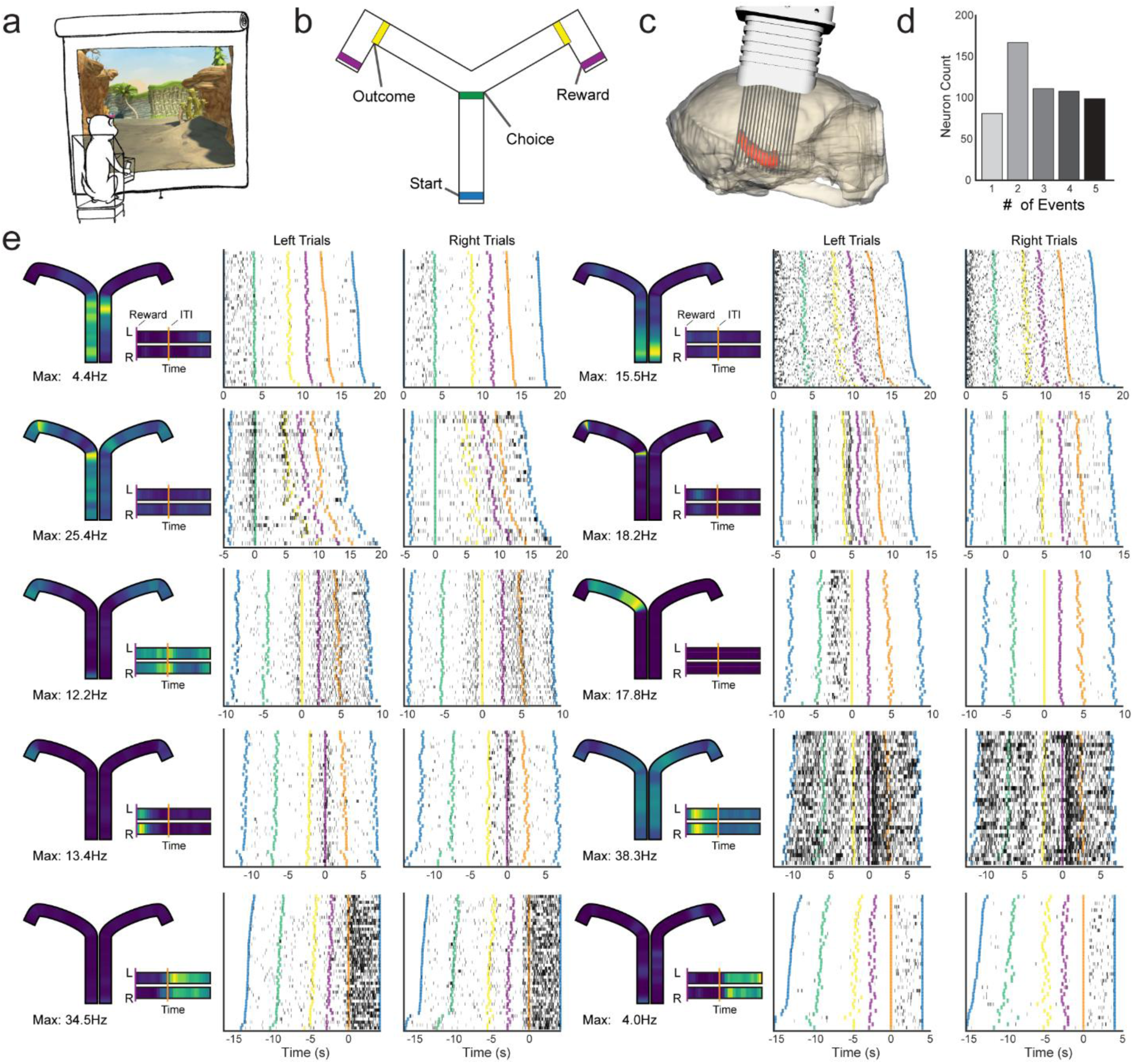
| Event-aligned responses of hippocampal neurons during a virtual alternation task. **a,** Immersive virtual Y-maze in which monkeys navigated via joystick and alternated between left and right arms across successive trials. Also see Supplementary Video 1. **b,** Top-down schematic of the maze illustrating the behavioral events used for alignment: movement initiation (Start, blue), turn initiation at the choice point (Choice, green), turn toward the reward (Outcome, yellow), reward delivery (Reward, purple). Not shown: onset of the inter-trial interval (ITI – black screen). **c,** Rendering of a chronically implanted microdrive targeting the hippocampus, based on MRI-derived 3D models of one subject. **d,** Distribution of event responsiveness across neurons. Histogram shows number of neurons that exhibit significant responses to multiple behavioral events, demonstrating that 485 neurons responded to multiple task events, whereas only 81 were selective for one event. **e,** Sample neurons illustrating diverse peri-event firing profiles aligned to the five behavioral events in **b**. Trials are ordered by duration for clarity. Colored lines indicate behavioral events as shown in **b**. For each neuron, leftward trials are plotted on the left and rightward trials are plotted on the right. Trials are aligned (time zero on the horizontal axis) to the behavioral event with the strongest alignment for each neuron. Maze schematics to the left of each set of peri-event plots show firing rates on left and right trials. Warmer colors indicate higher firing rates (color scale: 0 to max). Time insets to the right of each maze schematic show firing rates following reward onset (purple line) and ITI onset (orange line). The upper schematic in each inset corresponds to time following reward on leftward trials and the lower schematic corresponds to time following reward on rightward trials.

We recorded from the hippocampus of each monkey with a chronically implanted microdrive consisting of 124 independently movable single-wire electrodes (Fig. 1c). Recording locations were confirmed to sample the full longitudinal extent of the hippocampus through post-mortem histological reconstruction. To characterize how individual neurons responded to key features of the task, we aligned spiking activity to five behavioral events spanning all task phases: movement initiation from the central stem (Start), turn initiation at the choice point (Choice), turn initiation toward the reward location (Outcome), reward delivery (Reward), and onset of the inter-trial interval (ITI) (Fig. 1b). Individual neurons exhibited reliable responses aligned to these events (Fig. 1d,e), and the response profiles included both sharp modulation time-locked to specific moments and sustained but irregular epoch-spanning activity.

### Event-aligned responses across task phases

To systematically assess event-aligned responses, we analyzed activity within a 2-s window surrounding each event. We computed the mutual information between instantaneous firing rate and peri-event time for each neuron, with significance assessed against a within-trial circular-shift null distribution (*P*<0.01). Across the population of 1659 uniquely recorded neurons, 566 neurons showed significant modulation to at least one event, with responses distributed across spatial and non-spatial task phases: Start (n=342; 60.4%), Choice (n=305; 53.9%), Outcome (n=348; 61.5%), Reward (n=350; 61.8%), and ITI (n=329; 58.1%). Strikingly, the vast majority of these neurons (n=485; 85.7%) were significantly responsive to multiple events (Fig. 1d), indicating that these neurons do not act as feature detectors that are narrowly selective for specific task variables or sensory stimuli.

### Events segment the task into distinct epochs

We next explored how neuronal responses contribute to population-level representations across the task. Spikes were first binned in non-overlapping 50ms intervals to compute instantaneous firing rates over the course of each trial. To compare trials of variable duration, the timeline of each trial was rescaled piecewise between the onset of task events (Movement Start, Choice Turn, Choice Arm, Outcome Arm, Reward, and Inter-Trial Interval; Fig. 2a). That is, each inter-event interval was linearly stretched or compressed to match the mean duration for that epoch across the session, yielding a standardized trial lasting 17s that was sampled at 68 evenly spaced points (250ms resolution within rescaled time). Incorrect trials were excluded from further analysis. This temporal rescaling approach ensured that both movement periods and stationary epochs were represented (Fig. 2a). 588 neurons demonstrated significant mutual information (*P*<0.01) between instantaneous firing rate and the rescaled task timeline; these neurons composed the population used for subsequent analyses. We did not observe systematic differences in neuronal specificity along the hippocampal longitudinal axis or across subfields, so all hippocampal neurons were pooled for analysis (Supp Fig. 1).

**Figure 2.**
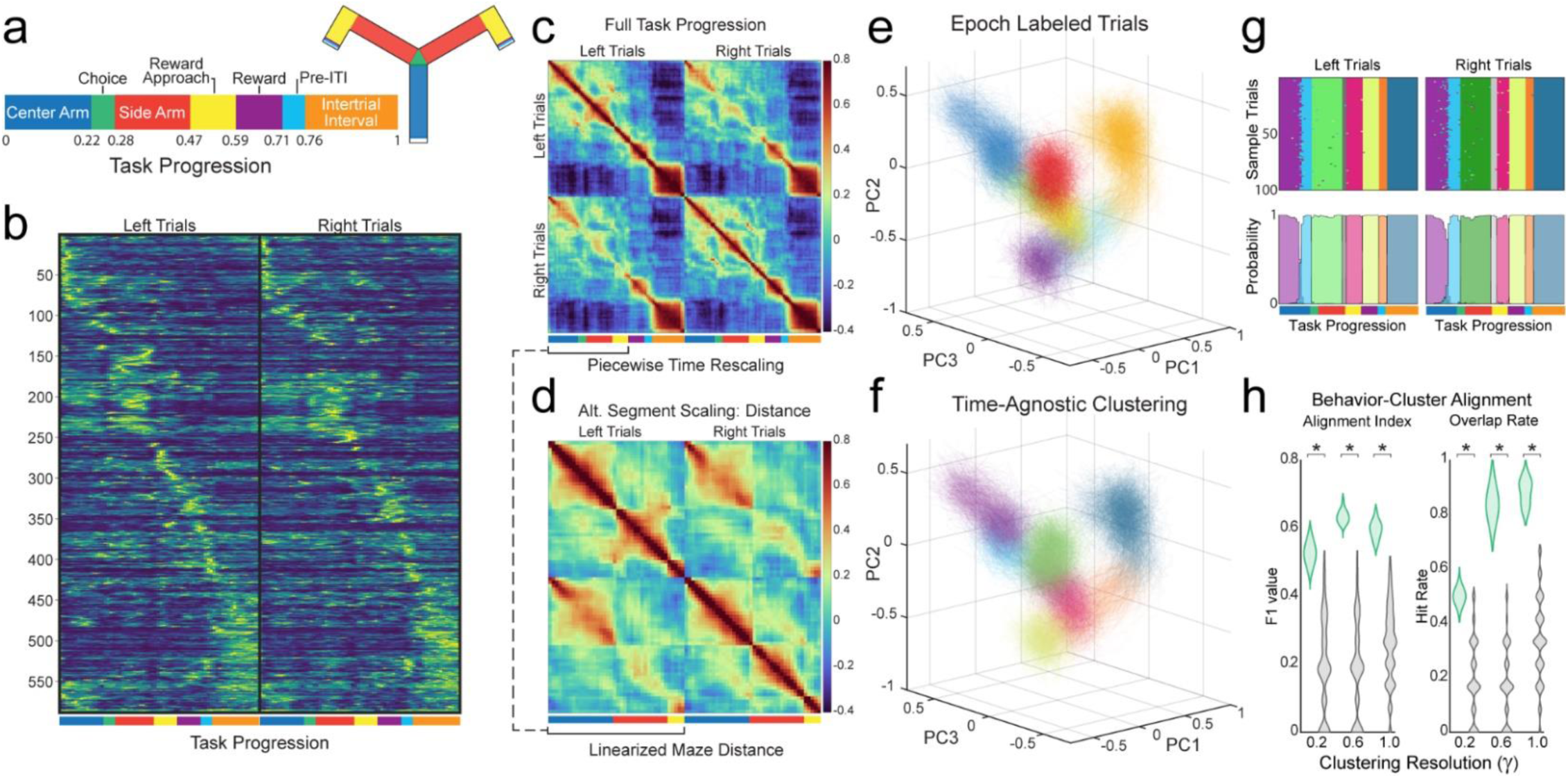
| Population activity segments the task into discrete event-aligned states. **a,** Standardized trial timeline generated by piecewise temporal rescaling between task events (left) with corresponding epochs mapped onto the 2D maze (right). **b,** Population heatmap showing mean firing rates (neurons × bins) across the standardized trial, revealing event-aligned structure. Warmer colors indicate higher firing rate (0 to max). **c,** Temporal cross-correlation matrix of soft-normalized population vectors, showing strong within-epoch similarity separated by sharp discontinuities at epoch transitions. **d,** Equivalent analysis to **c**, after rescaling trials by distance along the linearized maze, confirming consistent epochal segmentation under a spatial alignment framework. **e,** Three-dimensional PCA projection of pseudopopulation responses from 1,000 trials (left trials shown), color-coded by behavioral epoch as in **a**. **f,** Same projection as **e**, colored by behavior-agnostic cluster identities; clusters closely correspond to the behavioral epochs in **a**. **g,** Top: Cluster labels mapped back onto time for 100 left and 100 right pseudotrials, showing contiguous epoch-like blocks. Bottom: Probability of each cluster label across bins for left and right trials plotted as continuous curves, revealing reproducible boundaries aligned to behavioral events. **h,** Alignment of cluster-derived boundaries with behavioral transitions, quantified by F1 scores and Hit Rates relative to null distributions, demonstrating strong correspondence between unsupervised segmentation and task epochs. Asterisks denote significance (Mann–Whitney U tests, *P* < 0.001).

At the population level, these neuronal responses combined to form activity patterns that differentiated task phases across the trial. Rather than being uniformly dispersed across time, neuronal responses aligned to task events (Fig. 2b). Temporal cross-correlation of soft-normalized population vectors^68,69^ revealed sustained blocks of high within-epoch correlation separated by sharp discontinuities at event boundaries (Fig. 2c). These event-related discontinuities were not dependent on the specifics of the rescaling procedure: comparable block structure was observed when the navigation period was uniformly rescaled and with piecewise rescaling between alternative maze landmarks (Supp Fig. 2). Together, these results demonstrate that neural populations adopt distinct states within each task epoch, with transitions reliably aligned to behavioral events.

To verify these results were not dependent on temporal scaling, we rescaled trials by distance along the linearized maze. This approach similarly revealed within-epoch stability of the population response (Fig. 2d), confirming that hippocampal activity exhibits stable states within each phase of the task. While this spatial representation revealed within-epoch stability, it compresses temporally extended behavioral periods (e.g. turns) into single spatial coordinates. This binning increased off-diagonal correlations and emphasized the consistency within the long maze arms but obscured the fine temporal structure evident in event-aligned data. Overall, these analyses reveal that hippocampal population activity is organized into discrete epochs delimited by behaviorally salient events, a structure that reflects the temporal architecture of the task itself.

### Agnostic clustering of population activity matches behavioral epochs

To examine the variability of hippocampal dynamics on individual trials in light of this observed epoch structure, we analyzed pseudopopulations constructed from independently recorded neurons. Pseudopopulations provide a stringent test of whether behavioral epochs reflect generalizable organizational principles: if discrete clusters emerge from randomly sampled trials, it would indicate that the variability is sufficiently constrained to preserve distinct, epoch-specific population states. We generated large synthetic ensembles by sampling single trials from each neuron with replacement (1,000 left and 1,000 right pseudotrials), preserving each neuron’s temporal profile and trial-to-trial variability while achieving dense sampling across the recorded population (588 neurons). At each moment in the task, activity across all neurons defined a population vector capturing the instantaneous network activity. When projected into low-dimensional space, the vectors formed distinct clusters mirroring the behavioral epochs of the task (Fig. 2e – left trials only, Supp Fig. 3a – left and right trials, Supp Videos 2 and 3).

We applied unsupervised clustering to the pseudopopulation activity, entirely blind to behavioral labels. Each pseudotrial yielded 68 population vectors (one per timepoint), and all vectors were pooled across trials and timepoints before clustering (136,000 total vectors from 2,000 pseudotrials). Clustering was performed in the native 588-dimensional neural space using Pearson correlation as the similarity metric. This approach was agnostic to trial identity, timepoint, and behavioral labels (Fig. 2f –left trials only, Supp Fig. 3b – left and right trials, Supp Video 4), providing a data-driven method to assess whether latent structure aligns with behavioral segmentation.

Unsupervised clustering revealed that pseudopopulation vectors organized into discrete, highly reproducible clusters. We clustered the data using two independent algorithms, Leiden community detection and spherical k-means clustering, which both converged on highly similar partitions of the neural state space across parameter sweeps (Leiden: kNN ∈ {12, 24}, γ ∈ {0.2, 0.6, 1.0}; k-means: k ∈ {6–12}). Cluster assignments were consistent across repeated runs, with within-resolution Adjusted Rand Indices, a metric of similarity between clustering methods, exceeding null distributions generated from comparing circularly shifted pseudopopulations (Mann-Whitney U, all *P*<0.001, Supp Fig. 4f). Cross-parameter comparisons likewise yielded high Adjusted Rand Indices (Mann-Whitney U, all *P*<0.001, Supp Fig. 4f), indicating that similar cluster structure was preserved across parameter settings and that population organization was robust to changes in clustering resolution and algorithmic parameters. This convergence demonstrates that population vectors organize into discrete, stable clusters, revealing a structure that is intrinsic to hippocampal dynamics rather than an artifact of any particular clustering approach.

These clusters exhibited striking temporal organization. When mapped back onto the trial timeline, each cluster formed a nearly contiguous block rather than fragmenting randomly across time, thereby defining discrete epochs that were reproducibly expressed across trials (Fig. 2g). Cluster transitions were remarkably consistent across clustering approaches and replications, with boundaries between clusters aligning at the same positions in the trial timeline (Supp Fig. 4). These transition locations remained stable across all parameter sets and both clustering algorithms, even when the total number of clusters varied. While clustering resolution affected the number of clusters identified, higher resolutions split existing clusters rather than revealing new temporal boundaries, indicating that these transition points reflect fundamental organizing features of the population dynamics.

We tested whether these cluster-derived boundaries align with behavioral structure by comparing boundary timing to the events that delimit the behavioral epochs (Fig 2a). Focusing on boundary timing rather than cluster identity allowed us to assess alignment independent of clustering granularity, since boundaries remained stable even when cluster identities and cluster counts differed across resolutions. We quantified alignment using two complementary metrics: F1 scores, which balance the rate of correctly identified behavioral epoch transitions against false detections, and simple hit rates, the proportion of behavioral epoch transitions that overlapped with a cluster boundary. Across all parameter sets, true boundaries showed markedly higher alignment with behavioral epochs than null distributions (Fig. 2h; Mann-Whitney tests: all *P*<0.01). Mid-level resolutions produced clusters that nearly matched behavioral epochs in both number and transition timing, resulting in the highest precision (F1 scores) and near-ceiling accuracy (hit rates). Both higher and lower resolutions identified behavioral boundaries accurately but with reduced precision due to cluster count mismatches. Together, these results demonstrate that hippocampal population activity adopts discrete states that transition precisely at behaviorally relevant events—forming a structure robust enough to emerge from wholly unsupervised clustering.

### Variability across phases of the task reveals two modes of hippocampal processing

Having established epochal segmentation of hippocampal population activity, we next asked whether the reliability of population representations varies across the task. Specifically, we asked whether population activity is expressed more consistently at particular behavioral moments. We quantified this by correlating each trial’s activity against the leave-one-out population mean at each time bin, yielding a trial-by-template correlation matrix (Fig. 3a). Extracting the diagonal of this matrix—which represents the similarity between corresponding time bins—provided a continuous measure of trial-to-template similarity across the trial (Fig. 3b). This measure revealed that cross-trial similarity increased sharply at behaviorally–relevant events, indicating highly stereotyped population states, while the intervening epochs showed greater trial-to-trial variability. We constructed null distributions by disrupting temporal alignment through independent circular rotations of each neuron’s activity (1,000 iterations). True trial-to-template similarity significantly exceeded null expectations in restricted task windows (permutation tests, Benjamini-Hochberg corrected, *P* < 0.05), confirming that instances of high reliability reflected coordinated alignment of the population with task structure.

**Figure 3.**
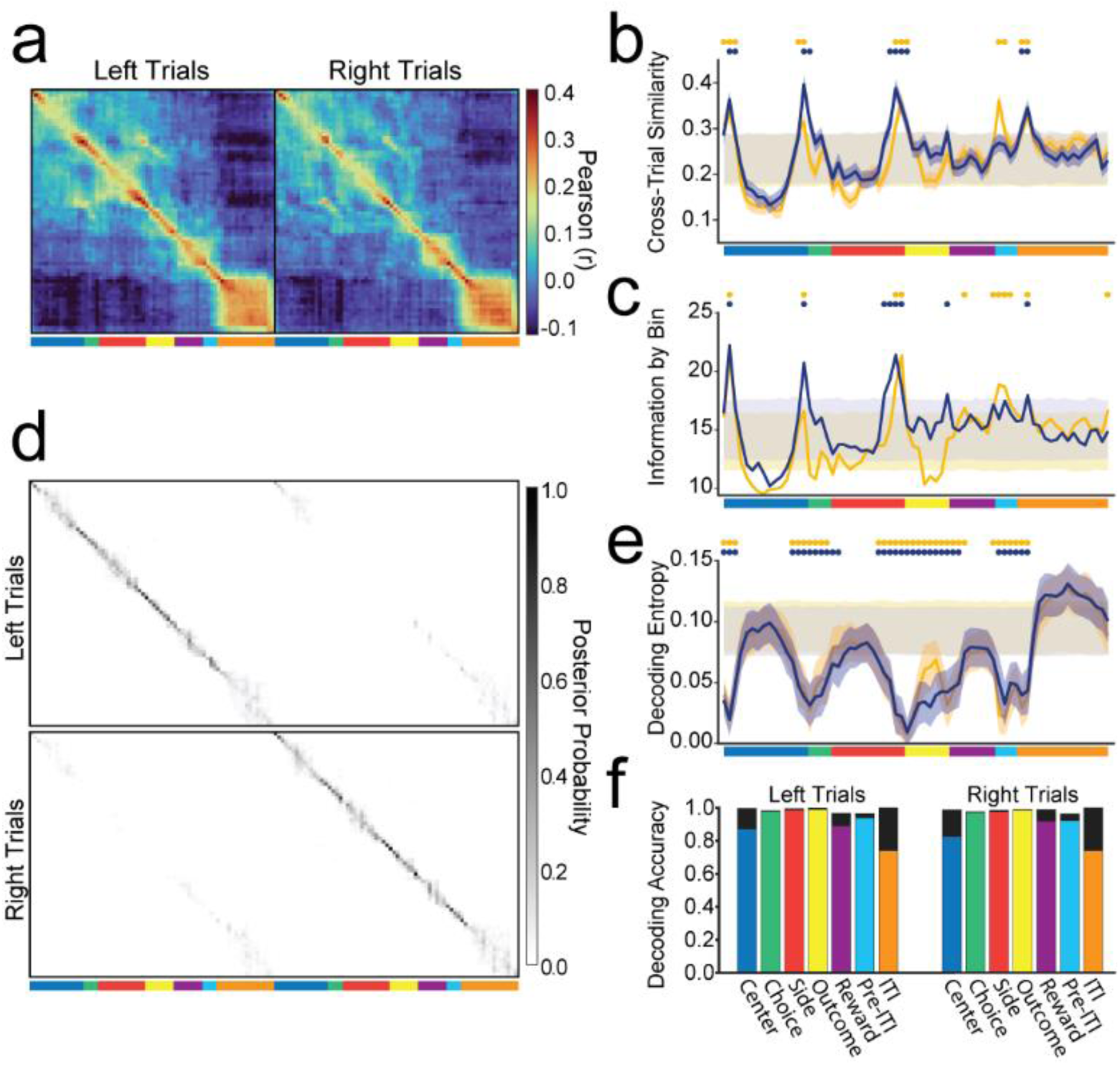
| Dual representational modes alternate across behavioral epochs. **a,** Mean temporal cross-correlation between each trial’s population vector and a leave-one-out mean template (averaged across trials) showing elevated cross-trial similarity at the behavioral events corresponding to epoch boundaries. **b,** Mean trial-to-template similarity (diagonal of **a**) across the standardized trial, showing peaks in cross-trial similarity at behaviorally-relevant events and reduced similarity within epochs. **c,** Aggregate information-by-bin for the population, revealing transient peaks in information content aligned to behavioral events. **d,** Bayesian decoding posteriors showing precise decoding at epoch boundaries and broader, but still epoch-specific, posterior distributions within epochs. **e,** Entropy of the posterior probability, quantifying decoding uncertainty, which is lowest at event boundaries and elevated within epochs. **f,** Epoch-level decoding accuracy across the standardized trial. Colored bars indicate decoding that is both epoch- and trial-type-specific, whereas black bars indicate decoding accurate for epoch identity irrespective of trial type. Dots denote bins significantly different from null (permutation test, *P*<0.01).

To further examine how reliably population activity patterns map onto temporal progression through the sequence of task events, we asked whether task progression could be decoded from the population activity across individual trials. Using a Bayesian decoder with a Poisson-based framework, we constructed templates from half of the trials and decoded the remaining half. For each time bin in a held-out trial, the decoder produced a posterior probability distribution over all time bins in the standardized trial, representing the likelihood that the observed population activity corresponded to each moment in the task.

Posterior probability distributions revealed differential precision across the task (Fig. 3d). At behaviorally salient events, posteriors were tightly concentrated around the true time bin, indicating precise temporal decoding. By contrast, during the epochs between events, posteriors were far more diffuse, spreading across multiple bins within the epoch. Remarkably, while posteriors were diffuse within epochs, they respected epoch boundaries: the decoder reliably identified *which* epoch was occurring, even when uncertain about *when* within an epoch. This pattern reveals that hippocampal population activity provides precise temporal information at behavioral transitions but represents moments within epochs categorically, reliably signaling epoch identity while being less informative about the exact moment within that epoch.

We further quantified these observations by measuring epoch-level decoding accuracy (Fig. 3f). Trial-type-specific decoding—requiring identification of both epoch and left/right trial type—showed high accuracy across epochs, although some epochs discriminated trial type better than others (all epochs >90% accurate, except ITI at 70%; χ² tests *P*<0.001 for each). When trial type was ignored, epoch identity was decoded with near-perfect accuracy (all >95% accurate; χ² tests *P*<0.001 for each). Thus, the population activity reliably signals the current phase of the task, irrespective of its ability to disambiguate how much time has passed within an epoch.

Beyond accuracy, posterior entropy provides a complementary measure that quantifies uncertainty of the decoded state (Fig. 3e). Entropy was lowest at event boundaries and elevated within epochs, quantitatively confirming that the decoder was highly certain at the times of events but uncertain about precise temporal position within epochs (permutation tests, Benjamini-Hochberg corrected, *P* < 0.05). This structure matched the pattern observed in the trial-to-template correlation analysis (Fig. 3b): both revealed high precision during behaviorally-relevant events, and elevated uncertainty during the intervening epochs, despite accurate decoding of epoch identity.

To probe the basis of this temporal structure, we next examined when individual neurons were most informative about task progression. We computed information-by-bin—a time-resolved estimate of mutual information between instantaneous firing rate and position in the task timeline^70^. Summing across all recorded neurons yielded the upper bound of the total information carried by the population at each time bin. To assess significance, we compared this to null distributions generated by independently circularly rotating each neuron’s trials, recalculating information-by-bin, and summing across the population. The population’s aggregate information showed prominent peaks aligned with behavioral events and epoch boundaries that significantly exceeded null expectations, with reduced information content in the intervening intervals (Fig. 3c). These information peaks mirrored the temporal structure revealed by the correlation and decoding analyses, with population activity most reliable at behavioral events and reduced information within intervening epochs (permutation tests, Benjamini–Hochberg corrected, *P*<0.05).

Across these three independent analytical frameworks—trial-to-template correlation, Bayesian decoding, and information content—a consistent pattern emerged. Hippocampal population activity exhibits two distinct representational modes that alternate across the task. At salient behavioral events, the population activity converges into precise, highly reliable states across trials that are informative about task progression. Between these events, activity is less reliable, exhibiting greater trial-to-trial variability and reduced temporal precision. Critically, this within-epoch variability does not reflect noise or decoupling from task structure: population activity remains confined to epoch-specific subspaces, reliably signaling which phase of the task is occurring even while being less informative about exact temporal position within that phase. This dual architecture points toward a fundamental organizing principle: the hippocampus does not represent experience as a continuous temporal stream, but rather constructs a temporal scaffold anchored to behaviorally relevant events with the intervals between these events represented holistically. This architecture may explain how the hippocampus balances the need for precise temporal ordering of events with the flexibility to generalize across variable durations and trajectories.

### Neuronal responses reflect abstract task structure

To test whether hippocampal activity reflects environment-specific features or the abstract structure of task phases, monkeys performed an A–B–A′ sequence of visually distinct but geometrically identical Y-mazes, with each maze run separated by >15 minutes of quiescence in dim lighting. The virtual environments were designed to maximize perceptual dissimilarity, with different landmark positions and stark contrasts in colors, surface textures, and decorations (Fig. 4a). Maze A′ repeated the visual appearance of maze A to assess within-session stability. We analyzed 306 neurons that were significantly responsive in maze A, maze B, or both across the full A–B–A′ regimen.

**Figure 4.**
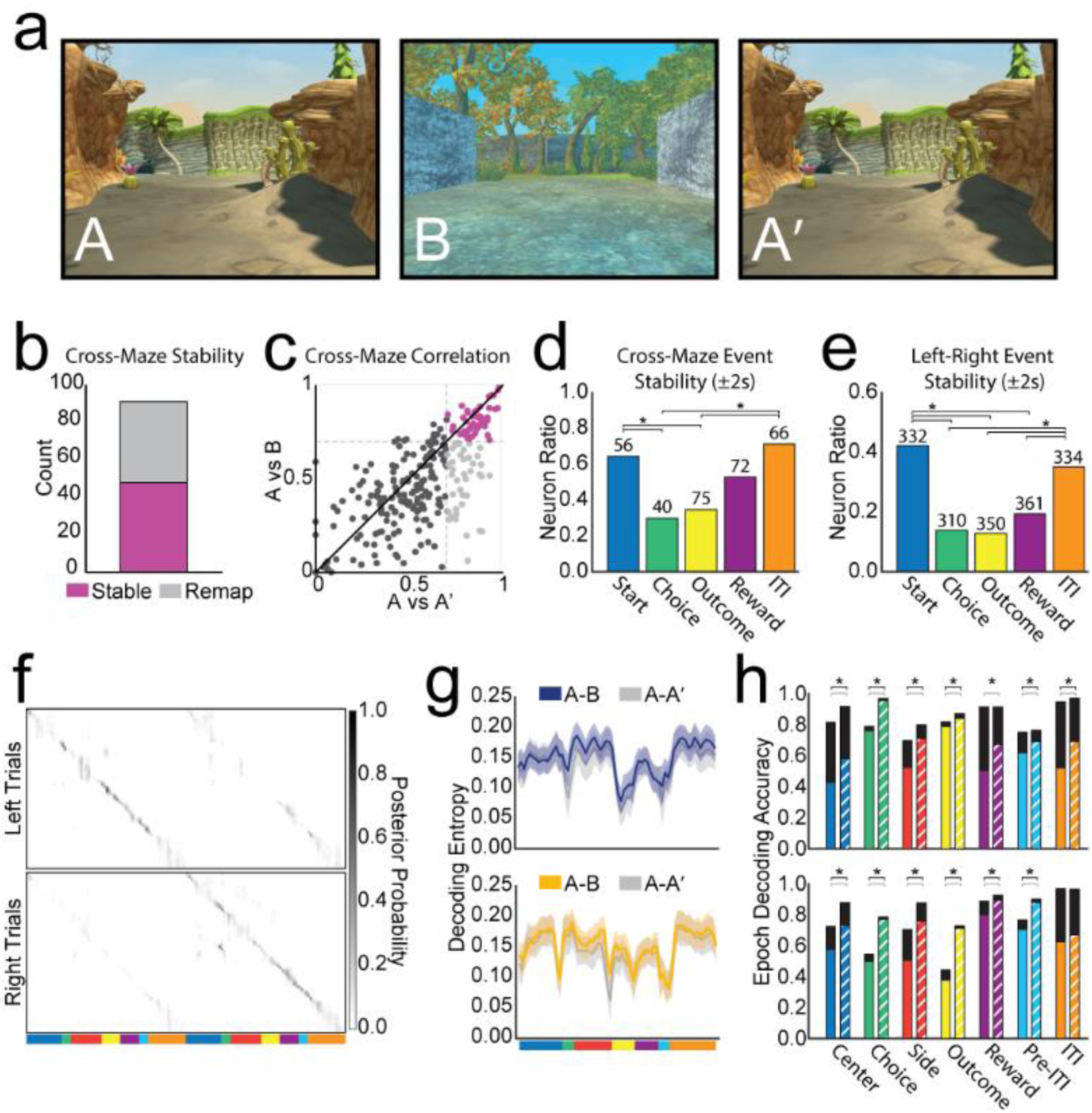
| Hippocampal representations generalize across visually distinct environments. **a,** Visually distinct virtual Y-mazes (A, B, A′) differing in landmarks, colors, and textures; maze A′ is a repeat of maze A. **b,** Counts of neurons classified as stable or remapped between mazes A and B, restricted to neurons stable between repeated presentations of maze A (A–A′). **c,** Correlation of full-trial firing patterns across mazes A–A′ (x-axis) and A–B (y-axis); dashed line indicates the stability threshold (r = 0.7). **d,** Event-wise stability across mazes A and B for neurons stable in A–A′, revealing differential preservation across task phases (asterisks denote *P*<0.001, χ² test). **e,** Event-wise stability across left and right trial types, illustrating elevated trial-type stability at Movement Start and the ITI initiation (asterisks denote *P*<0.001, χ² test). **f,** Cross-maze (A–B) Bayesian decoding posteriors for left (top) and right (bottom) trials, showing accurate tracking of task progression. **g,** Decoding entropy across the trial for A–B (color) and A–A′ (gray) conditions, plotted separately for left (top) and right (bottom) trials. **h,** Epoch-level decoding accuracy for A–B (solid color) and A–A′ (striped, same hue). Colored portions denote trial-type-specific decoding; black portions denote epoch-accurate decoding without regard for trial type (gray brackets indicate significant trial-type specific differences, and black brackets indicate significant differences irrespective of trial-type, *P*<0.01, χ² test). Top, left trials; bottom, right trials.

In rodent hippocampus, exposure to new environments typically triggers global remapping, where new place fields emerge and the active population changes^71^. By contrast, we found that many primate hippocampal neurons maintained stable responses across visually distinct mazes. We defined neurons as stable if they exhibited significant mutual information with task progression in both mazes (*P*<0.01) and Pearson correlations of trial-averaged firing rates greater than 0.7. Of the 91 neurons exhibiting stable responses between repeated presentations of maze A (A–A′), 50 (55%) preserved their response profiles between mazes A and B (Fig. 4b,c). The relatively low A–A′ stability (91/306) is consistent with the growing literature on representational drift, in which population codes evolve over time even in familiar environments^72,73^, here expressed on a rapid, multi-hour timescale^74^. Against this backdrop, the substantial preservation of responses across visually distinct environments (A-B) demonstrates that hippocampal activity can encode task structure independent of visual features.

Event-specific analyses revealed differential cross-maze stability across task phases. Restricting our analysis to neurons that were stable across A–A′, we assessed how many preserved their peri-event activity patterns (±2 seconds) when the environment changed (A–B). Stability was high at Movement start (64%) and ITI start (71%) but substantially lower at Choice (30%) and Outcome (35%) turns, while Reward start was intermediate (53%) (Fig. 4d; χ²(4)= 31.30, *P*<0.001). Post-hoc Fisher exact tests with Holm correction confirmed that Movement start and ITI start each differed significantly from Choice and Outcome events (adjusted *P*<0.05). These results indicate that the earliest and latest events of the trial carry the most task-generalizable coding, whereas the events in which the subject must attend to make a choice and evaluate the outcome are less likely to be stable across virtual environments.

We found a similar pattern of stability across left and right trials. Only 14% of responsive neurons were stable across left and right trials at the Choice turn and 13% at the Outcome turn, compared to substantially higher stability at Movement start (42%) and ITI start (35%) (Fig. 4e; χ²(4)=125.38, *P*<0.001). Post-hoc Fisher exact tests with Holm correction confirmed that Movement start and ITI start each differed significantly from Choice, Outcome, and Reward events (adjusted *P*<0.05). Together, these findings reveal a shared structure across maze and trial-type generalization: stable responses concentrate at the beginning and end of the trial, whereas decision and feedback epochs exhibit more situational specificity where attention to task contingencies is necessary to be successful.

Population-level analyses confirmed that hippocampal representations generalize across visual environments. A Bayesian decoder trained on maze A accurately tracked task progression when applied to maze B, with posterior probability patterns and entropy closely matching within-environment decoding (A′ from A) (Fig. 4f). Decoding entropy across the trial was indistinguishable between A→B and A→A′, indicating that overall uncertainty was not affected by environment identity (Wilcoxon rank-sum test with Benjamini-Hochberg correction, *P*>0.05 all bins; Fig. 4g). The characteristic two-mode structure (i.e. Fig 3d) persisted: posteriors were sharply concentrated at event boundaries and more diffuse within epochs, with diffusion constrained by epoch identity. For most epochs, epoch-level decoding accuracy was higher during repeated presentations of the same maze than during presentations of a maze with a different appearance (χ² tests p<0.01; Fig. 4h), reflecting environment-specific responses in a subset of neurons. Nonetheless, decoding epochs of maze B from the maze A template significantly exceeded chance (χ² tests *P*<0.001 for each; chance 5–24% proportional to epoch durations). Although decoding accuracy was modestly lower for A→B, epoch-level decoding was conserved across environments, indicating that primate hippocampal populations can encode the ordinal structure of task events in a manner that abstracts over sensory context. Together, the convergence of single-neuron stability and population-level decoding demonstrates that a substantial portion of the hippocampal population responds to the abstract structure of the task rather than to the specific visual environment.

## Discussion

The hippocampus is central to memory and cognition, but the organizing principles of its activity have remained elusive. Here, we show that hippocampal population dynamics are organized not by individual sensory or spatial features but by the temporal structure of the engaged task. Individual neurons responded to multiple behavioral events, indicating that they do not act as narrow feature detectors but participate in a distributed representation spanning the full task sequence. These population responses segmented the task into distinct phases, each defined by a unique ensemble code. Transitions were precisely aligned to behavioral events, revealing that the hippocampus partitions experience into meaningful elements. Many neurons exhibited similar activity patterns across visually distinct environments, showing that the underlying code generalizes beyond sensory context^75^ to capture the abstract progression of task events. Together, these results reveal that hippocampal activity reflects the learned temporal organization of behavior: a structure that persists across contexts and provides a framework for linking elements within an episode.

A central and novel finding of this study is that hippocampal population activity alternates between two distinct modes across the unfolding of experience. At behaviorally-relevant events, hippocampal activity converged into tightly coordinated, low-variability configurations that were highly reproducible across trials. Between these events, activity patterns were more variable, dispersing within epoch-specific regions of population state space. One possible explanation for this pattern is that externally driven input at behaviorally-salient moments triggers transitions between different intrinsically constrained states. This view aligns with the broader cortical observation that salient input acutely quenches variable ongoing neural activity, producing reliable population responses and a reproducible post-event state^76,77^. During the inter-event intervals, variable activity may be governed by loosely constrained intrinsic dynamics within each epoch-specific subpopulation. These dual modes suggest that the hippocampus toggles between moments of input-driven “transitions” and intrinsically organized “maintenance” states, thereby stitching a discontinuous series of goal-directed events into a continuous episode. This dynamic could implement a temporal chunking mechanism in which extrinsic input creates anchors in time while internally sustained activity bridges the intervals between them—together forming a scaffold for organizing experience. The ability to link events across time may offer a unifying explanation for the fundamental contribution of the hippocampus to memory and cognition.

These dynamics offer a new lens through which to interpret the functional role of the hippocampus. Traditional accounts have largely focused on the content of hippocampal representations, often casting the hippocampus as a conjunctive feature detector^26,27^ or part of a circuit specialized for performing metric spatial calculations^4,5^. More recent frameworks emphasize the process by which these representations evolve over time to structure experience^28,29,34^. The present findings build on this perspective and suggest the intriguing hypothesis that hippocampal population activity is organized by transitions between externally anchored and intrinsically sustained states, a dynamic architecture that links discrete behavioral events into a coherent temporal progression. This organization resonates with theories of event segmentation and predictive processing: the hippocampus not only encodes the elements of experience but also tracks the ordinal relationships among them^34,78^. Individual events serve as nodes in a graph-like scaffold connected by bridging activity that captures their temporal relationships. By projecting the complex multidimensionality of experience onto a compact cognitive graph^79,80^, the hippocampus provides a framework for organizing behavior and enabling inference across time. The capacity to bind separated events into unified representations may underlie the hippocampus’s broad role in memory and cognition^32,81^—establishing the computational foundation for linking episodes as coherent elements of experience.

## Acknowledgements

This work was supported by the NIH (BRAIN Initiative – U19NS107609 (EAB, ALF); NIMH – R01MH117777 (EAB, JWR); NINDS – UF1NS126485 (EAB, ALF); NIDA – 5T90DA032436 (YB); ORIP – P51OD010425), the McKnight Endowment Fund for Neuroscience (EAB), the Simons Foundation, Simons Collaboration for the Global Brain (EAB, ALF), and the Wayne E. Crill Endowment at the University of Washington (EAB). We are grateful to Megan Jutras for outstanding project support, Dr. Sierra Schleufer and Ian O’Leary for excellent technical assistance, and Drs. Charles Gray and Baldwin Goodell for microdrive development. We also thank Drs. Larry Squire, Lara Rangel, Andrew Alexander, and Gregory Horwitz for insightful discussions.

## Author Contributions

J.W.R., Y.B., A.L.F. and E.A.B. conceived the study. J.W.R., Y.B. and E.A.B. designed the experimental methods. J.W.R. and E.A.B. performed surgical procedures. J.W.R., Y.B., A.J.M. and B.K. carried out behavioral training, electrophysiological recordings and histological reconstruction. J.W.R., A.J.M., Y.B. and B.K. performed spike sorting, unit curation and data preprocessing. J.W.R. and Y.B. performed single-neuron analyses. J.W.R. performed population-level analyses. J.W.R. prepared the visualizations. J.W.R. and E.A.B. wrote the original manuscript draft. J.W.R., Y.B., A.J.M., B.K., A.L.F. and E.A.B. edited and revised the manuscript. E.A.B. supervised the project. J.W.R., Y.B., A.L.F. and E.A.B. acquired funding.

## Supplementary Information

Supplementary Information is available for this paper, including videos demonstrating behavior and data visualizations.

Correspondence and requests for materials should be addressed to Jon W. Rueckemann or Elizabeth A. Buffalo.

## Methods

### Animals

All experiments were carried out in accordance with National Institutes of Health guidelines and were approved by the University of Washington Institutional Animal Care and Use Committee. Three adult male rhesus monkeys (*Macaca mulatta*; M1: 9.3 kg, 7.5 years old; M2: 9.1 kg, 8.5 years old; M3: 12.3 kg, 8 years old) were used in the experiments.

### Y-maze behavioural task

Monkeys were trained to manipulate a joystick mounted to their protective chair. Subjects first learned to center and approach a visual cue displayed on an immersive screen to obtain reward. After mastering joystick control, animals were trained to navigate simple linear paths in an immersive virtual environment. Virtual environments and task mechanics were developed in the Unity game engine.

Monkeys then performed a spatial alternation task in a Y-shaped virtual maze (Fig 1, Supp Video 1). The maze comprised three arms converging at a central choice point. Subjects navigated a semi-constrained path via joystick, which prevented backward movement or lateral deviation except at designated turn points.

Each trial began with the subject positioned at the base of the central stem. After traversing the stem, the subject reached a junction where it could select the left or right side arm. Reward contingencies required alternation between arms across successive trials, such that the correct choice on a given trial depended on remembering the outcome of the previous trial. At the end of a correct arm, subjects rounded a second corner to encounter a visual cue (a banana) marking the reward location. The cue was occluded until the corner was reached, ensuring that the outcome remained unknown until the side arm had been fully traversed. Contact with the banana triggered delivery of a slurry of banana and monkey chow (Purina).

On correct trials, reward was available for ∼2s, followed by 1s of immobility before the screen went black for a 4s inter-trial interval (ITI). On incorrect trials, subjects were prevented from rounding the final corner, and the ITI was initiated immediately upon reaching the end of the chosen arm. Following the ITI, the subject was repositioned at the start of the central stem for the next trial. After initial training and during recording sessions, monkeys rarely made errors, performing consistently at greater than 95% accuracy.

Ten visually-distinct virtual environments were created for this experiment, and monkeys experienced multiple mazes in an A-B-A design, allowing for the determination of environment-specific responses in the neurons (i.e. remapping). Each recording session, monkeys experienced one or two of the ten visually-distinct environments.

The events used to calibrate peri-event spiking activity (Fig 1) were (i) movement initiation on the center stem (Start), (ii) turn initiation at the choice point (Choice), (iii) turn initiation toward reward location (Outcome), (iv) reward delivery (Reward), and (v) onset of the inter-trial interval (ITI).

### Implantation of chronic microdrive

A titanium headpost (Gray Matter Research) was implanted near the posterior vertex of the skull. After acclimation and task training, each monkey was implanted with a 124-channel microdrive (LS124; Gray Matter Research) targeting the full longitudinal axis of the right hippocampus. Chambers and drives were custom fit to each animal’s skull and dural surface using preoperative MRI, ensuring electrode trajectories were directed toward hippocampus. Each channel carried an independently-movable tungsten electrode (100μm diameter, ∼1 MΩ impedance; FHC) with more than 40 mm travel distance from the bottom of the microdrive. Implantation followed a three-stage procedure: 1) the chamber was implanted and sealed; 2) after healing and verification of sterility, a craniotomy was performed and sealed with a form-fitting plug; and 3) the sterile, custom microdrive was secured within the chamber and hermetically sealed.

Electrodes were advanced to ∼10 mm below the brain surface on the day of implantation, then lowered over the next week to 2–3 mm above target. Each electrode was advanced via a fine-threaded screw (125μm per revolution).

### Electrode positioning

During sedation, a sub-millimeter 3D surface scan of each animal’s head and implant was obtained (Spyder; Artec). The surface mesh was registered with the preoperative MRI (Slicer 5.6.1) to ensure accurate alignment of the chamber relative to hippocampal anatomy and to generate precise trajectory projections for each electrode. Electrode depth was tracked through detailed turn records, allowing estimation of recording target positions. Adjustments were guided by audio monitoring of multi-unit spiking activity and visual inspection of local field potentials, which helped account for potential deviations in electrode trajectory. Electrodes were advanced only after completion of a recording session, allowing them to settle for at least a day before the next session to mitigate within-session drift. Not every electrode was moved each day; instead, advancing a subset was often sufficient to shift surrounding electrodes into fruitful recording positions.

### Histological reconstruction

At the conclusion of experiments, 30μA of direct current was passed through each electrode to produce small electrolytic lesions at the electrode tips. After a minimum two-day survival period to allow gliosis, animals were transcardially perfused with 4% paraformaldehyde with electrodes in place. Following 48 hours of post-fixation, the microdrive was carefully withdrawn. The skull was positioned in a stereotax, and the brain was blocked in the coronal plane to facilitate matching of histological sections to the preoperative MRI. Tissue was cryoprotected by graded infiltration of glycerol and DMSO, before being rapidly frozen in −70°C isopentane^82^. Brains were coronally sectioned at 50μm, and every third section (150μm spacing) was Nissl-stained with cresyl violet.

A 10x micrograph was acquired for each Nissl section. Custom software used an iterative alignment procedure to register each section both to the preoperative MRI and to adjacent sections, resulting in a 3D Nissl volume stitched from the micrographs and coregistered to the MRI. A unique rescaling factor (1.75–2.50%, empirically determined for each animal) was applied to compensate for tissue shrinkage during cryoprotection.

The 3D Nissl volume and projected trajectories from the 3D surface scan were used to identify each electrode track in the tissue. This alignment enabled full reconstruction of the true path taken by each electrode to its terminal lesion. Daily recording positions were estimated from turn records as the relative distance along the electrode tract from the electrolytic lesion site. For electrodes that followed a straight trajectory, premortem position estimates were typically accurate within ∼250μm; however, a notable subset showed deviations due to bending.

### Electrophysiological recording

Neural signals were recorded with a Neuralynx Digital Lynx SX system (Neuralynx Inc., USA) using 124 channels sampled at 30 kHz with 24-bit A/D conversion. Broadband signals were acquired and grounded to the titanium recording chamber, with all channels referenced to ground.

Spike detection was performed offline. Continuous data were first high-pass filtered using a 4th-order Butterworth filter at 400Hz (zero-phase, ‘filtfilt’ MATLAB). Candidate spikes were identified by threshold crossing at 3.5 standard deviations below the mean of the filtered trace.

### Spike sorting

Candidate units were clustered automatically with Isosplit (MountainSort^83^). Clusters were then triaged using custom visualization software that projected the spikes onto peak–valley and principal-component axes. This tool enabled rapid identification of (i) MountainSort-identified clusters consistent with noise for exclusion and (ii) channels without MountainSort clusters that contain putative units for rescue.

All putative clusters were subsequently curated manually in Offline Sorter (Plexon, USA). Sorting decisions were based on cluster density and separation in feature space. Units were accepted if they formed a stable and distinct cluster with consistent waveform morphology; clusters showing a lack of separation, infiltration by noise, or significant drift over the course of recording were excluded.

### Temporal standardization of trials

To enable comparison across trials of variable durations, each trial was normalized to a standardized timeline sampled at 68 evenly spaced predictor bins. Because only the movement portion of the task varied in duration, four rescaling schemes were evaluated for this interval. The primary approach divided trials based on six task events (Movement Start, Choice Turn, Choice Arm Entrance, Outcome Arm Turn, Reward Start, and Inter-Trial Interval Start), with each epoch linearly rescaled to match its mean duration across sessions, yielding a 17-second trial with 250-ms bin resolution (Fig 2). Three alternative schemes served as controls: (1) uniform rescaling across the entire movement period, (2) piecewise rescaling between alternative landmarks (Movement Start, Center Stem Midpoint, Side Arm Midpoint, Reward Start, and Inter-Trial Interval Start), and (3) nonlinear rescaling based on distance in the linearized maze rather than elapsed time. Results throughout use the primary piecewise method; all schemes yielded consistent findings (Fig 2, Supp Fig 2).

Spikes were first binned at 50-ms resolution to preserve fine temporal structure. These 50-ms rates were then mapped to the nearest predictor bin. For trial rate maps, samples falling within the same predictor bin were averaged. This yielded one value per predictor bin per trial for population analyses. For mutual information calculations, individual 50-ms samples were retained to capture within-bin variability.

### Neuron inclusion criteria

Units were considered eligible for analysis if they fired at least 100 spikes within at least one maze during a given session, provided that maze also contained ≥50 correct trials. In addition, the unit was required to fire at least once on ≥25% of those trials for inclusion.

Neurons were classified as task responsive if they showed both elevated information content and consistent trial-level activity patterns. For each neuron, trial activity was represented as binned spiking across the full trial timeline (predictor details above). Three statistics were computed: (i) Shannon mutual information between binned spikes and trial progression, (ii) correlation of trial-averaged activity between the first and second halves of trials, and (iii) correlation between odd- and even-numbered trials of the same trial type (left–left, right–right). Significance for all three measures was determined relative to a null distribution (1000 iterations) generated by independently circularly shifting spikes across time within each trial. A neuron was classified as task-responsive if its observed statistic exceeded the 99th percentile of this null.

To mitigate the possibility of counting the same neuron across days, multiple criteria were applied to define unit uniqueness. A unit was considered unique if (i) the electrode from which it was recorded had been advanced, (ii) more than 5 days had elapsed since the last included recording on that channel, or (iii) the Pearson correlation of its trial-averaged activity with that of the previous day was <0.5. In addition, neurons were conservatively excluded if their mean activity subjectively appeared similar to nearby recordings during manual curation.

### Shannon mutual information

Mutual information between instantaneous firing rate and bin identity was computed from all 50-ms rate samples. Let *r* denote the discretized firing rate and *x* the predictor bin (68 x 2 trial-type). Information was given by:

The marginal and conditional probabilities were estimated as *P*(*r*)and *P*(*r* ∣ *x*), and information was given by:

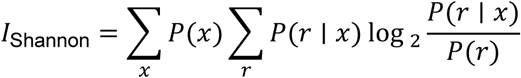

where P(x) is the probability of occupying bin x, P(r|x) is the conditional probability of observing rate r in bin x, and P(r) is the probability of observing rate r. Spike counts were quantized in two-spike increments to keep response categories compact in high-firing neurons, maintaining a manageable dynamic range and ensuring comparability of information estimates across neurons.

Significance was assessed against a within-trial circular-shift null (1000 iterations) in which each neuron’s spike train was independently circularly shifted within each trial.

### Skaggs information

Information per spike was computed according to Skaggs and colleagues^84^, using the trial-average rate curve, r(x) obtained by averaging 50-ms rate samples within each predictor bin. Trial-average rate curves were Gaussian smoothed with a kernel of 3 bins.

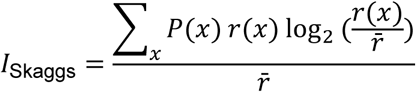

where ***r̅*** = ∑_*x*_ *P*(*x*) *r*(*x*). This normalization expresses information in bits per spike, facilitating comparison across neurons with different overall firing rates.

### Sparsity

Tuning compactness of the smoothed trial-averaged rate curves r(x) was quantified by probability-weighted sparsity^85^:

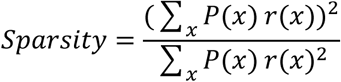

### Population analysis

Population activity across the standardized trial is visualized as an NxB matrix of mean firing rates, where N is the number of neurons and B is the number of predictor bins (B=136; 68 bins per trial x 2 trial types). For the population rate plot (Fig 2b), firing rates for each neuron were averaged across all correct trials and linearly rescaled from 0 to 1 by dividing by the neuron’s maximum rate. For visualization, neurons were grouped by the similarity of their mean activity profiles using hierarchical clustering, and the resulting clusters were reordered to reflect the general temporal progression of activity across the trial.

#### Rate soft-normalization

To balance the contribution of neurons with differing overall firing rates in subsequent population-level analyses, each neuron’s binned activity was soft-normalized^68,69^. For each neuron, the full set of predictor-binned firing rates—concatenated across all trials—was treated as a single time series. The transformed rate was given by:

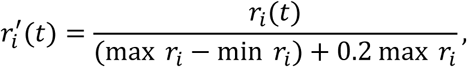

where *r*_*i*_(*t*)denotes the binned firing rate of neuron *i* at time *t*.

This transformation preserves each neuron’s relative modulation while reducing disparities in scale across the population. Soft-normalized firing rates were used for temporal cross-correlations and agnostic clustering, but were not used in Bayesian decoding.

#### Temporal cross-correlations

To quantify the temporal organization of ensemble activity, we computed pairwise correlations between population vectors across all predictor bins. For each bin (B), the population vector comprised the soft-normalized firing rates of all neurons. The resulting BxB matrix captured the similarity of ensemble activity patterns across trial progression, with blocks of high correlation indicating stable population states and sharp boundaries marking event-aligned transitions.

### Behavior agnostic clustering

To identify latent structure in population dynamics independent of task labels, we applied unsupervised clustering to neural activity patterns without incorporating behavioral epoch boundaries or trial-type information. Because neurons were recorded across different sessions, we constructed pseudopopulations that combined neurons while preserving each cell’s temporal dynamics and trial-to-trial variability.

#### Pseudopopulation construction

For each trial type, we generated synthetic population trials (“pseudotrials”) by randomly sampling, with replacement, one real trial from each task-responsive neuron (n = 588), using their binned, soft-normalized firing rates. The selected trials were then combined such that activity across all neurons formed a single population vector at each time bin. Each pseudotrial therefore consisted of a sequence of 68 population vectors corresponding to 68 time bins that span the full standardized trial. This procedure was repeated 1,000 times per trial type, producing a large set of pseudotrials that densely samples the distribution of network activity while preserving each neuron’s temporal response profile and natural trial-to-trial variability.

#### Null distributions

To assess statistical significance of clustering structure, null pseudopopulations were created by disrupting temporal coordination across neurons while preserving each neuron’s firing rate distribution. For each neuron, the binned rate time series was circularly shifted by a random offset independently across trials, eliminating temporal alignment across the population but maintaining within-trial firing statistics. Pseudotrials were then assembled from these permuted data using the same sampling procedure, repeated 1,000 times to produce null ensembles matched in size and composition to the empirical sets. Test statistics from empirical pseudopopulations were compared against distributions generated from these null ensembles.

#### Clusters methods

Population vectors from all time bins and trials were pooled and clustered directly in high-dimensional space using Pearson correlation distance. Dimensionality reduction via PCA was used only for visualization, not for clustering. Two complementary unsupervised methods were applied:

##### Leiden community detection

The Leiden algorithm^86^ identifies clusters by optimizing community structure in a k-nearest-neighbor (kNN) graph. We constructed kNN graphs^87^ (Facebook AI Similarity Search (FAISS), Meta AI Research) with k = 12 (corresponding to the connectivity threshold ln(n) for the dataset size) and k = 24 (as a higher-connectivity comparator). For each graph, clustering used the Reichardt–Bornholdt configuration model with three resolution parameters (γ = 0.2, 0.6, 1.0), which control the propensity to split communities. Each parameter combination was replicated 100 times to assess partition stability.

##### Spherical k-Means clustering

As an alternative approach, spherical k-means clustering was applied with correlation distance for k = 6–12 clusters. Each fit was optimized across 100 random initializations, retaining the solution with the lowest within-cluster sum of squares. This process was repeated ten times for each value of k to determine consistency.

#### Cluster stability assessment

Clustering consistency across clustering runs was quantified using Adjusted Rand Index (ARI). The Rand Index (RI) measures the proportion of bin pairs on which two labelings agree:

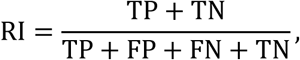

where TP and TN are correctly grouped or separated pairs, and FP and FN are errors. The Adjusted Rand Index (ARI) corrects for chance agreement,

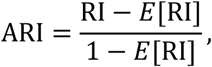

where *E*[RI]is the expected RI under random labeling.

For each parameter set (e.g., Leiden: kNN = 12, γ = 0.6), metrics were computed across all pairs of replicate runs to assess within-parameter stability. Metrics were also compared across different parameter sets to assess consistency of boundary placement across connectivity and resolution choices.

Metrics were computed both directly (unmatched) and after Hungarian matching of cluster identities. Unmatched comparisons penalize both differences in cluster assignments and differences in the number of recovered clusters. Matched comparisons compensate for hierarchical splits or merges and isolate whether partitions preserve consistent boundary placement irrespective of cluster count. This dual approach distinguished instability in the number of clusters from instability in boundary placement, allowing focus on temporal precision of state transitions rather than categorical labels.

### Cluster boundary detection

To isolate the temporal precision of population state transitions from differences in cluster granularity, we focused analyses on when cluster identity changed (corresponding to presumptive population state changes) rather than on the cluster identity itself. This boundary-based approach asks when transitions happen, independent of what the cluster labels are or how many clusters are recovered. The high trial-to-trial consistency of clustering outcomes enabled meaningful comparison of state transition timing across trials and clustering runs (Fig 2g).

#### Template construction

For each clustering run, we identified the most common cluster label at each time bin across trials. These modal labels defined a cluster template representing the dominant segmentation pattern for that run. The template thus captured the typical sequence of population states without requiring that every trial follow this exact pattern.

#### Boundary identification on individual trials

Trial-specific boundaries were identified as time bins where the cluster label changed relative to the preceding bin. To exclude brief, single-bin fluctuations within otherwise stable epochs, a boundary was considered legitimate only if the transition occurred within ±3 bins of the corresponding template boundary. This criterion ensured that detected boundaries represented sustained state transitions consistent with the majority pattern across trials.

#### Aggregation across trials and runs

For each clustering run, the probability of observing a legitimate boundary at each bin was calculated across trials, yielding a boundary likelihood profile. These profiles were then aggregated across the 100 replicate runs within each parameter set (e.g., Leiden: kNN = 12, γ = 0.6), producing a run-by-bin matrix of boundary likelihoods (Supp Fig 4). This matrix served as the foundation for analyses of boundary prevalence and behavioral alignment. Averaging this matrix yielded a boundary prevalence curve (values 0–1) representing the probability of a boundary at each bin for that parameterization. Peaks in the prevalence curve indicated bins where boundaries consistently recurred across replicates.

Equivalent prevalence curves were computed for null pseudopopulations generated by independent circular shifts of neuronal activity.

#### Statistical significance

To assess significance, we quantified the probability that null boundaries occurred within ±1 bin of each true prevalence peak. Multiple comparisons were controlled using the Benjamini–Hochberg procedure, identifying bins where observed boundary recurrence exceeded null expectations. These boundary prevalence curves were subsequently used to assess alignment with behavioral epoch boundaries.

#### Clustering alignment to behavioral events

Comparisons were made on boundary locations rather than cluster labels to test whether population transitions aligned with behavioral transitions independent of cluster count or identity. To test alignment with behavior, cluster-derived boundaries were compared to behavioral epoch boundaries using ±1 bin tolerance. The F1 score was used to quantify overlap accuracy across clustering resolutions by accounting for the true and false hit rates. True positives (TP) were cluster boundaries within ±1 bins of an epoch boundary; false positives (FP) were boundaries outside this window; and false negatives (FN) were epoch boundaries without a nearby cluster boundary.

Precision and recall were calculated as

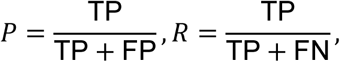

and combined as

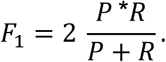

Distributions of F1 scores from true vs. null clusterings were compared with Mann–Whitney tests.

### Trial-specific analyses

#### Trial-wise correlation

To quantify cross-trial consistency, we compared each pseudotrial’s binned population vector to templates derived from leave-one-out (LOO) means. Separate templates were constructed for left and right trials by averaging across all other trials, excluding the current one. For each trial, the Pearson correlation was computed between its activity vector at each bin and every bin of the templates, yielding a cross-temporal similarity matrix. Rows of this matrix capture the similarity of each moment in the trial to the entire template trajectory, providing a measure of both moment-to-moment alignment with the expected template and cross-temporal structure that reflects variability in representational entropy.

To summarize alignment over time, we extracted the diagonal of the cross-temporal matrix, representing the similarity between each trial bin and the corresponding bin of the template. These diagonal values were averaged across trials to yield a mean similarity curve, with confidence intervals obtained by repeating the pseudopopulation sampling procedure 1,000 times.

Null distributions were generated by independently circularly rotating each neuron’s activity across bins, preserving marginal firing statistics while disrupting coordinated temporal structure. The LOO procedure was repeated 1,000 times on these null populations, producing null similarity matrices and curves. Statistical significance was assessed binwise by comparing each point on the true curve to the null distribution via permutation tests, with multiple comparisons controlled using the false discovery rate (q < 0.05).

#### Bayesian decoding

Neuronal population vectors were decoded under a Bayesian framework with a Poisson firing model. Pseudopopulation trials were split 50/50: templates were constructed from the mean firing rate of each neuron at each bin in the training set, and the test set was decoded by computing posterior probabilities over bins.

The posterior probability of bin *b* given observed spike counts **n** = (*n*_1_, *n*_2_, …, *n*_*N*_) from N neurons was computed via Bayes’ rule:

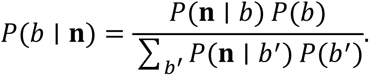

Here *P*(*b*)is the prior probability (assumed uniform across bins), and *P*(**n** ∣ *b*) is the likelihood under a Poisson model:

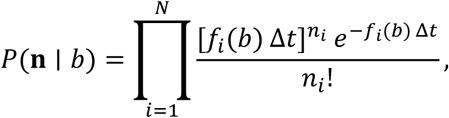

where *f*_*i*_(*b*) is the mean firing rate of neuron *i* in bin *b* (from the training template) and Δ*t* is the bin duration.

This cross-validation was repeated 1,000 times with random splits to construct confidence intervals for the entropy and accuracy quantifications below.

#### Decoding uncertainty

Entropy of the posterior probability across trials was used to assess decoding uncertainty. Entropy at each task progression bin was quantified as:

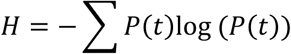

where *t* indexes the bin of population template (templates for left and right trials are concatenated). The mean entropy curve across cross-validations was used for statistical comparison to the null distribution via permutation tests at each bin, with multiple comparisons controlled using the false discovery rate (q < 0.05).

#### Decoding accuracy

Decoding accuracy was defined as the cumulative posterior probability assigned to the correct behavioral epoch. This border of the behavioral epoch was given a ±1 bin tolerance to account for slight timing jitter (∼250ms). Accuracy was computed both with trial-type separation (left vs. right) and after collapsing across trial types to account for trial-type specificity versus generalization.

Chance baselines were defined by the fraction of bins belonging to each epoch relative to total template length, adjusted for tolerance windows. Bin-wise accuracies were compared against chance using exact Wilcoxon signed-rank tests (right-tailed), with rank-biserial correlation as effect size. In complementary analyses, mean epoch accuracy was tested relative to chance with a binomial test. These approaches produced consistent results.

#### Information-by-Bin

To localize how information content varied across the standardized trial structure, we computed information-by-bin, a time-resolved adaptation of the “information-by-position” metric introduced by Olypher and Fenton^70^. Spike counts were quantized in two-spike increments to keep response categories compact in high-firing neurons.

For each neuron, the probability distribution of discretized spike counts was estimated within each predictor bin (*x*) and compared to the overall spike-count distribution across all bins. The resulting information-by-bin value,

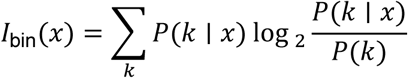

indicates how much the spiking statistics in bin x differ from those observed overall—i.e., how informative that bin of the trial is about the neuron’s firing pattern.

For completeness, the total mutual information for the neuron is the weighted sum of these binwise contributions,

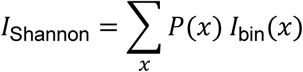

which expresses the overall reduction in uncertainty about trial time provided by the neuron’s firing activity (calculated above).

To estimate the reliability of neuronal activity at the population level, information-by-bin values were summed across all significantly responsive neurons. A comparator null was constructed by independently circularly shifting the information-by-bin curve for each neuron and then summing these shifted curves across the population. This procedure was repeated 1,000 times to generate a null distribution. Statistical significance was assessed binwise by comparing each point on the true curve to the null distribution using permutation tests, with multiple comparisons controlled using the false discovery rate (q < 0.05).

### Cross-maze analyses

#### Cross-maze consistency and decoding analysis

Neuronal activity was recorded as animals performed an A–B–A′ sequence of visually distinct but geometrically identical virtual mazes. Maze A′ repeated the original visual appearance to provide a control for within-session stability. To be considered for analysis, a neuron must be significantly responsive in one of the mazes, and each of the other mazes must contain ≥50 correct trials, have at least 100 spikes and the unit was required to fire at least once on ≥25% of trials on each maze. A total of 306 of the 588 significantly-responsive neurons were recorded under the full A-B-A’.

#### Individual neuron analysis

For each neuron, the average firing rate profile across the full trial timeline was compared pairwise across mazes (A–B, A–A′). A neuron’s response was classified as ‘stable’ between two mazes if: (1) it exhibited significant Shannon information in both mazes (as defined in the neuron inclusion criteria above), and (2) the Pearson correlation of averaged firing rates between mazes was >0.7.

For event-specific analyses, peri-event responses were quantified in a ±2 second window centered on each task event (Movement Start, Choice Turn, Outcome Turn, Reward onset, ITI onset). Spike counts were binned at 50-ms resolution to calculate mutual information across peri-event time bins. These binned rates were averaged across trials within each maze to generate peri-event rate vectors. The same stability criteria used for full-trial responses (significant mutual information in both mazes; Pearson correlation of peri-event rate vectors >0.7) were applied to classify each neuron as stable or unstable at each event.

To assess whether stability generalized across trial type rather than across maze appearance, the identical peri-event procedure was repeated for left vs. right trials within the same maze. Event-specific stability rates were therefore computed both for maze identity generalization (A–B) and trial-type generalization (left–right).

Differences in event-wise stability proportions across the five events were evaluated with χ² tests (df=4). Planned post-hoc pairwise comparisons were performed using Fisher exact tests with Holm correction for multiple comparisons.

#### Population decoding

A Bayesian decoder trained on maze A activity was used to decode trial progression in maze B and maze A′ pseudopopulations. Posterior probability matrices were compared across conditions. Similar to within-maze Bayesian decoding above, precision was quantified as the entropy of the posterior distribution, and accuracy as the likelihood of correctly decoding the behavioral epoch.

Decoding uncertainty (posterior entropy) was compared *per time bin* between cross-maze conditions (A→B vs. A→A′) using Wilcoxon rank-sum tests, with Benjamini–Hochberg false discovery rate control across bins (p=0.01).

Epoch-level decoding accuracy was evaluated in two parallel ways: (1) accuracy defined relative to the correct epoch *within trial-type* (left and right analyzed separately), and (2) accuracy defined relative to the correct epoch *after collapsing across trial types* to assess structure that generalizes beyond lateralization. For each definition, accuracy proportions were compared between cross-maze conditions (A→B vs. A→A′) using χ² tests conducted independently within each epoch. Chance baselines were determined by the fraction of template bins belonging to each epoch (±1 bin tolerance), and accuracy for each condition was compared against chance using χ² tests. Multiple epoch-wise comparisons were corrected using Holm adjustment.

## Data availability

The data that support the findings of this study are available from the corresponding author upon reasonable request.

## Code availability

The custom code used to analyze the findings of this study are available from the corresponding author upon reasonable request.

## Extended Figures and Supplemental Material

**Supplemental Fig. 1.**
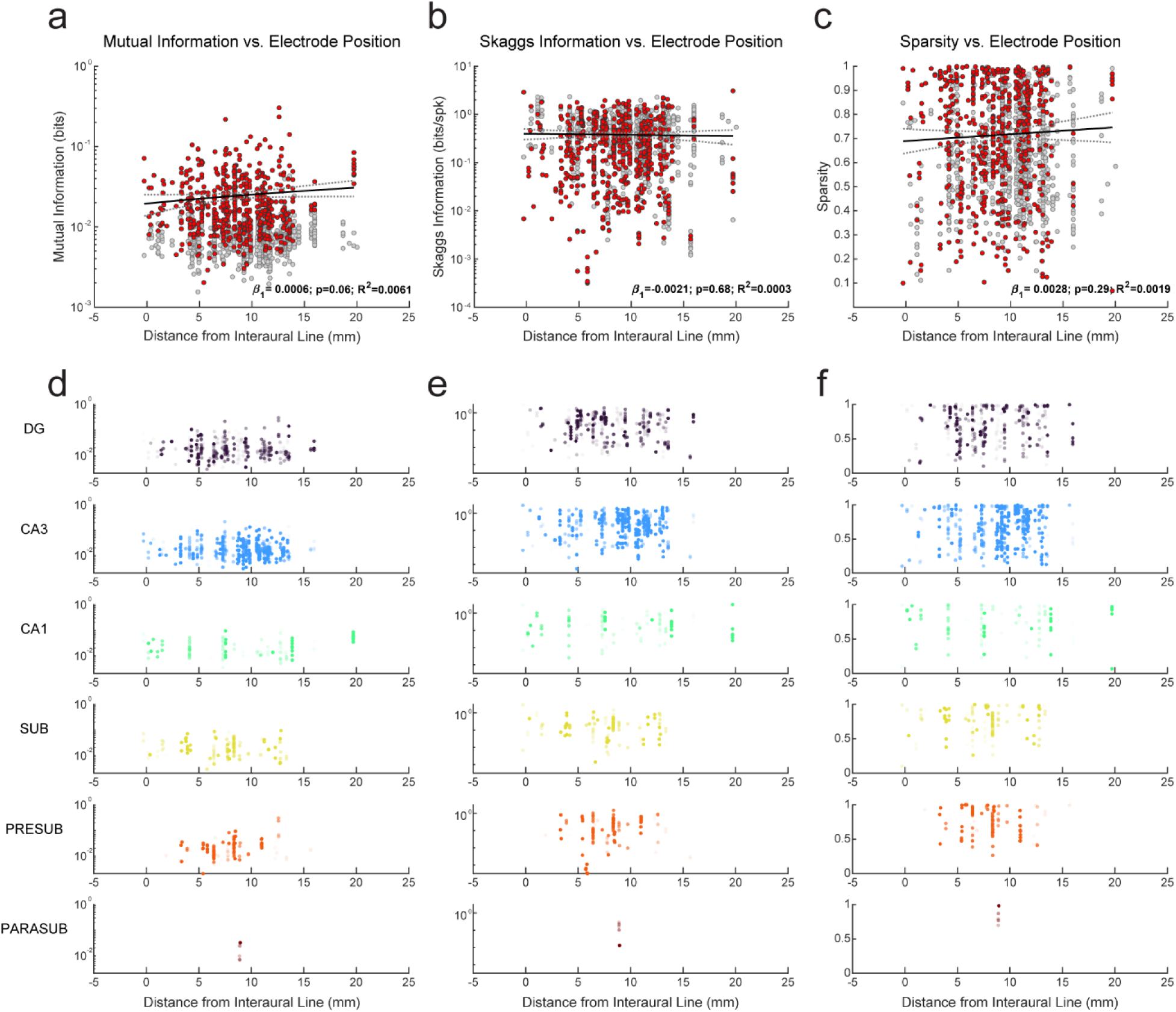
| Neuronal spiking specificity along the anteroposterior axis of the hippocampus. **a–c**, Relationship between neuronal specificity and distance from the interaural line (mm) across all recorded neurons (n = 588). Shown are mutual information (**a**), Skaggs information (**b**), and sparsity (**c**). Neurons with significant mutual information (exceeding the 99th percentile of within-trial circular-shift null distributions) are shown in red; remaining neurons are shown in gray. Black lines indicate linear regression fits with dotted lines denoting 95% confidence intervals. Mutual information and Skaggs information are displayed on a logarithmic axis for visualization; regression statistics were computed on untransformed values. Reported β coefficients, P values (two-sided t-test on regression slope), and R² values are from the linear model. **d–f**, Same information metrics shown separately for hippocampal subregions: dentate gyrus (DG), CA3, CA1, subiculum (SUB), presubiculum (PRESUB), and parasubiculum (PARASUB). All neurons are plotted in each subregion panel; dot opacity reflects each neuron’s assignment probability to the indicated subregion, quantified as the fraction of labeled histological voxels within 1000 μm of the reconstructed recording location that belong to that subregion.

**Supplementary Figure 2.**
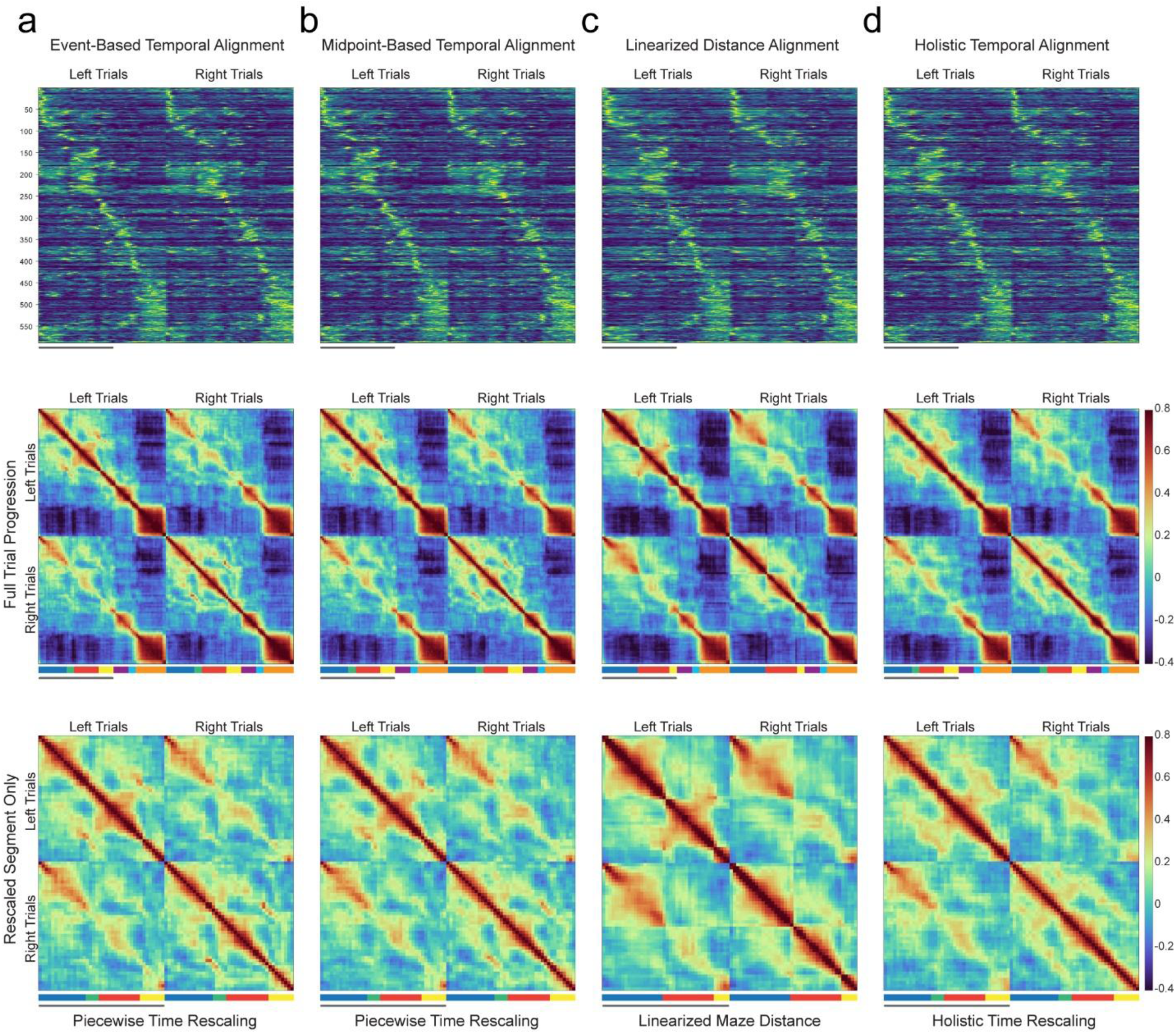
| Epoch structure is preserved across alternative alignment schemes. **a-d,** Columns compare four trial-alignment approaches: **a**, event-based temporal rescaling, **b**, midpoint-based temporal rescaling, **c**, linearized-distance alignment, and **d**, holistic temporal rescaling. All analyses use the same 588 task-responsive neurons. Top row: population heatmaps (neurons × bins) showing mean firing rates for each alignment method (0 max). Middle row (Full Trial Progression): Temporal cross-correlation matrices of soft-normalized population vectors, revealing sustained within-epoch blocks under all four alignment methods. Bottom row (Rescaled Segment Only): Enlarged view of cross-correlation matrices restricted to the trial segment warped by each alignment method, demonstrating that the pattern of state transitions is robust to the specific warping scheme used.

**Supplementary Figure 3.**
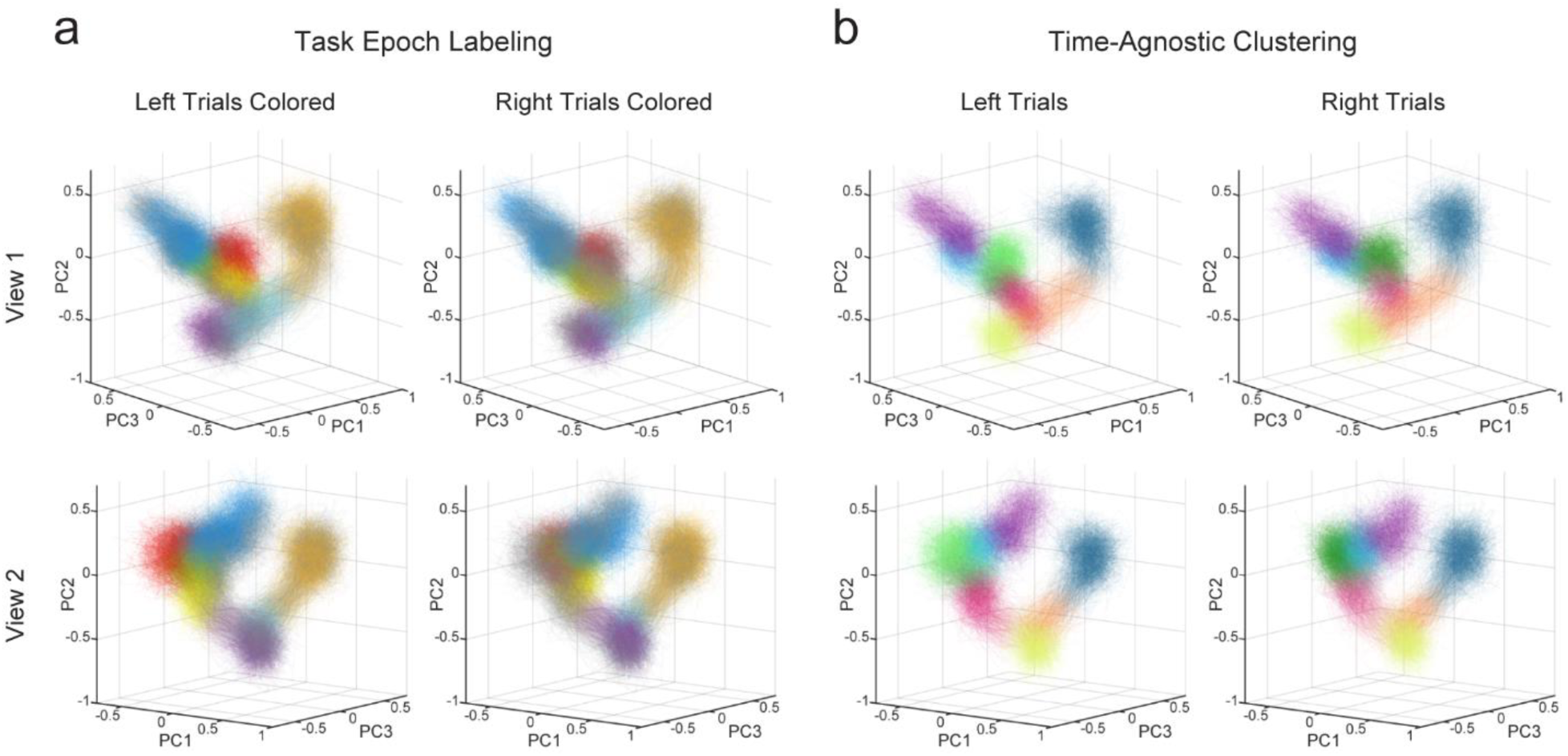
| Population state-space trajectories and behavior-agnostic clustering for left and right trials. **a**, Three-dimensional PCA projections of pseudopopulation responses from 500 pseudotrials, shown separately for left and right trials. Neural trajectories are color-coded by behavioral epoch, as in the main text. Left column shows left trials colored; right column shows right trials colored. Top and bottom rows show the same data from two different viewing angles (bottom row rotated 90° relative to the top). Similar to Fig 2e. **b**, Same projections as in **a**, colored by behavior-agnostic cluster identities derived from unsupervised clustering. Left column shows left trials only; right column shows right trials only. Cluster identities and colors are identical across left and right trials, except for clusters corresponding to side-arm traversal and turns toward the outcome arm (light/dark green and dark/light gray clusters, respectively). Viewing angles match those in **a**. Similar to Fig 2f.

**Supplementary Figure 4.**
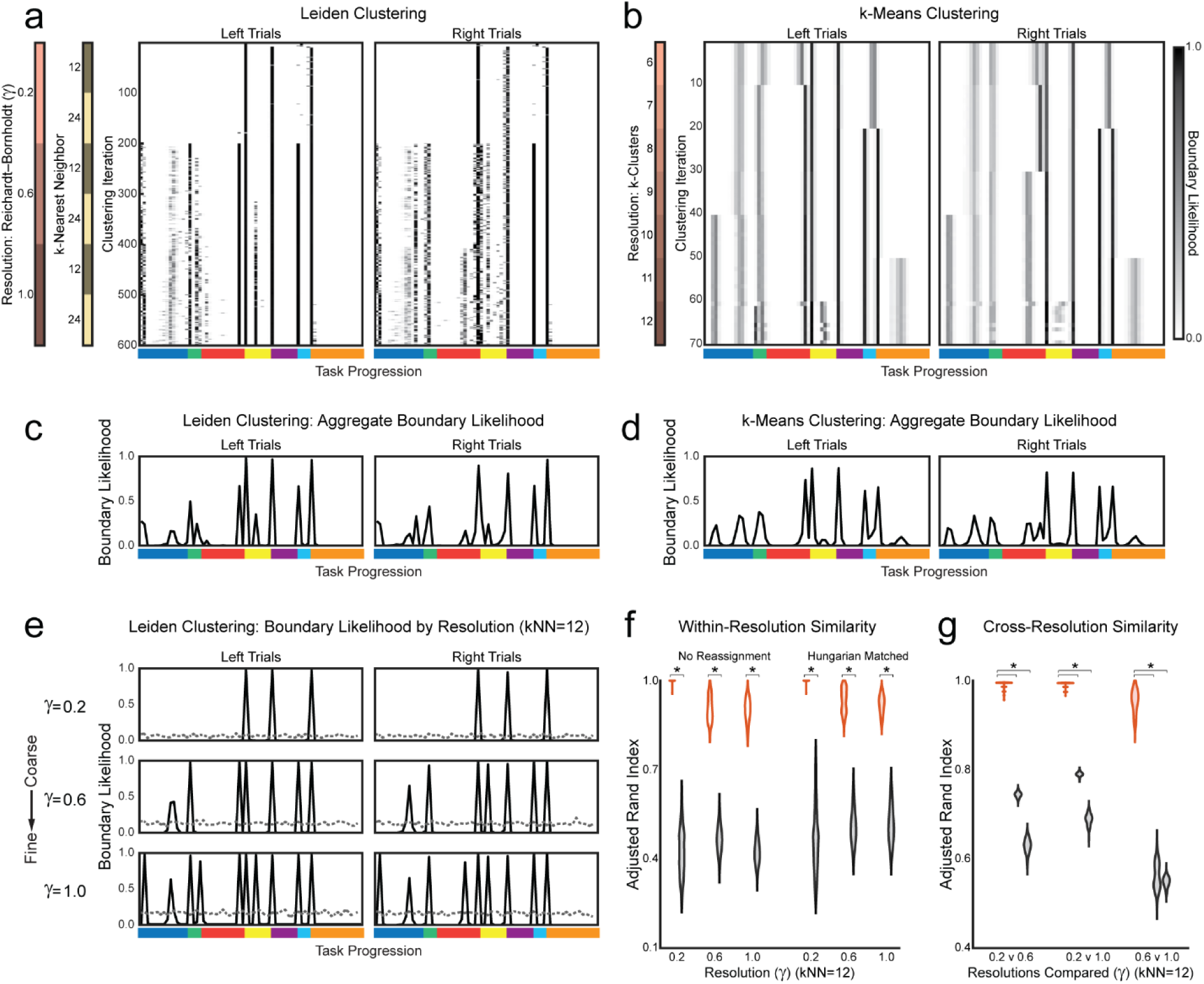
| Convergent boundary structure across clustering algorithms and parameterizations. **a**, Boundary likelihood for Leiden clustering as a function of task progression. Each row reflects one clustering run (100 replicates per parameter set), ordered by resolution γ ∈ {0.2, 0.6, 1.0} and kNN ∈ {12, 24}. Boundary likelihood reflects the proportion of trials with a cluster transition at each bin. **b**, Same analysis as **a** for k-means clustering (k = 6–12; 100 replicate initializations per k). **c**, Aggregate boundary prevalence curve for Leiden clustering (across all resolutions and kNN parameters), showing consistent peaks at common temporal locations. **d**, Aggregate boundary prevalence curve for k-means clustering. **e**, Boundary prevalence curves for kNN = 12 Leiden clustering shown separately for coarse, intermediate, and fine resolutions (γ = 0.2, 0.6, 1.0). Dotted lines indicate the distribution of peak likelihood across 1000 null pseudopopulations (circularly shifted pseudotrials). **f**, Within-resolution similarity of cluster solutions for Leiden clustering (kNN = 12). Violin plots show pairwise Adjusted Rand Index (ARI) across replicate runs for true and null pseudopopulations, for γ = 0.2, 0.6, and 1.0. Left set of plots shows ARI without compensating for cluster-count mismatches; right set shows ARI after greedy Hungarian matching of cluster identities. Significance bars reflect Mann–Whitney U tests, *P* < 0.001. **g**, Cross-resolution similarity for Leiden clustering comparing γ = 0.2 vs 0.6, 0.2 vs 1.0, and 0.6 vs 1.0. Violin plots show ARI after greedy Hungarian matching; each comparison includes one true distribution and two null distributions (directional asymmetry accounted for by swapping true and null label sets). Significance bars reflect Mann–Whitney U tests, *P* < 0.001.

**Supplemental Video 1. First-person view of the immersive virtual alternation task.** The monkey controls movement through the environment using a joystick. Two elements were added post hoc: an inset in the lower left showing the monkey manipulating the joystick, and a blue circle surrounding a white dot indicating the projected gaze position on the screen. In each trial, the monkey navigates along the central stem to a choice point and must alternate between selecting the right and left arms to obtain reward. Reward and reinforcement beeps are delivered upon collision with the banana object, followed by a 4-s intertrial interval during which the screen remains black until the next trial begins. Clicking sounds reflect the activity of a single neuron that fires at the choice point and during the turn toward the reward.

**Supplemental Video 2. Mean population trajectory and structured variability across task epochs of individual pseudotrials.** Three-dimensional PCA projection of hippocampal population activity, showing the mean population trajectory constructed by averaging across left trials for each of the 588 reliably task-responsive neurons. The trajectory is colored by task epoch and progresses continuously through the trial. This is followed by the progression of 10 population pseudotrial trajectories, each constructed by sampling one trial per neuron. Variability across pseudotrials gives rise to a diffuse cloud of non-parallel trajectories within each epoch. Despite this dispersion, trajectories converge at epoch boundaries coincident with salient task events, followed by rapid transitions into the next distinct subregion of state space.

**Supplemental Video 3. Task-epoch structured hippocampal population trajectories.** Three-dimensional PCA projection of hippocampal population activity, showing 500 population pseudotrial trajectories. Each pseudotrial samples one trial per neuron from the 588 reliably task-responsive neurons. Trajectories are colored by task epoch. Left-trial trajectories are shown first in color, then fade to gray as right-trial trajectories are introduced. Right trials are colored using the same epoch scheme. After completion of the right trials, left trials regain their epoch colors to visualize both trial types together. Within each epoch, population activity is constrained to a compact cluster within a subregion of state space, whereas transitions between epochs are marked by brief convergence events that shift the population trajectory into the next distinct subregion.

**Supplemental Video 4. Behavior-agnostic clustering of population state space trajectories.** Three-dimensional projection of hippocampal population activity onto the first three principal components, showing 500 left population pseudotrial trajectories. Each pseudotrial samples one trial per neuron from the 588 reliably task-responsive neurons. The full trajectories are initially shown in gray, and the view is rotated by 90° to emphasize their geometric structure. They are then color-coded according to cluster labels obtained using Leiden community detection (kNN = 12, γ = 0.6; same parameters as in Fig. 2f). The axes are then rotated back by 90° to return to the original viewing orientation, enabling direct visualization of the cluster organization within the population state space.

